# Learned Geometry, Predicted Binding: Structurally-Based Prediction of Peptide:MHC Binding Using AlphaFold 3 Enables CD4 T Cell Epitope Prediction

**DOI:** 10.1101/2025.07.10.664203

**Authors:** Kamel Lahouel, Mete Mulazimoglu, Jorge Soria-Bustos, Kameron Bates, Erin J Kelley, Lawson J Woods, Kunjur Manasa Upadhyaya, Gonzalo R Acevedo, Sophie Pénisson, Matteo Munini, Ehsan Variani, Margaret E Feeney, John A Altin, Cristian Tomasetti

**Author notes:** First Co-author.

## Abstract

The accurate prediction of T cell epitope peptides within proteins of interest has a wide range of applications, but is complicated by the multiple determinants of antigenicity, the polymorphism of the Major Histocompatibility Complex (MHC) locus, and the great diversity of possible peptide antigens. Leading in silico methods use a variety of statistical approaches to learn from sequences identified through both in vitro MHC binding and peptide elution studies, but their performance remains imperfect, particularly for MHC II-restricted responses. Here we present MHCIIFold-GNN, an entirely orthogonal solution to this problem that combines three new elements: (i) a highly-multiplexed peptide:MHCII binding assay, (ii) generalizable structural modeling using AlphaFold3, and (iii) transfer learning with a Graph-based Neural Network. Trained exclusively on newly-generated in vitro binders, we show that MHCIIFold-GNN enables state-of-the-art prediction of CD4 T cell epitopes presented by diverse MHC II proteins, on par with non-structural methods that rely on much larger datasets including naturally-processed ligands. Moreover, when MHCIIFold-GNN and a leading non-structural method are combined, we observe unparalleled performance on a held-out test set (11 % boost), underscoring the orthogonality of the methods. These results highlight the power of a new class of structure-informed approaches to the CD4 T cell epitope prediction problem.

## 1 Introduction

CD4 T cells are central mediators in immune responses to pathogens, tumor antigens, allergens and autoantigens, with effector functions triggered by the highly-specific recognition of peptide antigens [5, 18]. An essential step in this process is the binding of peptides to MHC [2, 29], a polymorphic and promiscuous family of proteins that serves the imperative of making peptides maximally visible to T cells, both within and across individuals [13]. Nonetheless, only a small fraction of possible peptides is capable of raising a T cell response in the context of a given MHC. While MHC binding is not the only requirement for T cell antigenicity, the accurate and efficient identification of this subset of peptides - which is both MHC- and antigen-specific - is a necessary and critical step in the development of research tools, diagnostics, therapies and vaccines in a wide range of areas [31, 23, 9]. However, the extensive population polymorphism of the MHC system as well as the great diversity of possible peptides within any given antigen target make this task extremely challenging. Early studies used in vitro molecular binding assays or elution of natural ligands to characterize the binders of individual MHC proteins, revealing allele-specific binding motifs and providing the foundation for initial models capable of predicting the binding potential of uncharacterized peptides to particular MHCs [14, 11, 30, 7, 34]. Initially, these approaches used relatively simple statistical methods trained on hundreds to thousands of data points, and showed modest predictive accuracy [21]. Subsequently, peptide:MHC datasets have grown in size and richness, powered largely by the increased throughput and sensitivity of mass spectrometric detection of naturally-presented ligands eluted from MHC-genotyped cells [27]. These developments have enabled the construction of more sophisticated models for the prediction of MHCII-restricted epitopes based on neural networks – including NetMHCIIpan [22], MixMHC2pred [22, 25] and MARIA [4] – and also enabled extrapolation to uncharacterized MHCs. These models generally operate on linear amino acid sequences as input features and use either fully connected neural networks [10] or architectures with more biological inductive bias, such as recurrent neural networks (RNNs) [10], to learn peptide:MHC binding preferences. Despite these advances, the accuracies of MHC class II models are substantially lower than class I models, especially for the T cell epitope prediction measure. This is likely in part due to lower resolution of peptide binding registers resulting from the open-ended nature of the MHCII binding groove. Most recently, breakthroughs in sequence-to-structure protein modeling using deep learning and exemplified by AlphaFold [33, 15] have generated growing interest in augmenting peptide:MHC prediction models by including three-dimensional structural modeling. It has been shown that AlphaFold can be used to accurately predict peptide:MHC structures21, and that a fine-tuned version for peptide binding prediction can enable prediction of MHC binding status with accuracy approaching that of the state-of-the-art [20]. AlphaFold 3 (AF3) is a recently-developed model that improves on earlier versions, notably by simplifying the way that aligned sequences are represented and introducing a generative diffusion step to refine the predicted atomic coordinates [1]. Previous AlphaFold models demonstrated the ability to infer accurate atomic-level structures from amino acid sequences. AF3 shows less reliance on sequence alignment and far superior performance on protein complexes, including antibody-antigen complexes – properties that we reasoned could also enable enhanced modeling of the highly-diverse peptide:MHC interface and thereby of T cell antigenicity. Another key advancement in AF3, from our point of view, lies in its ability to produce relevant intermediate and auxiliary feature representations of the input complexes. Therefore, it can be viewed as a foundation model, trained on large datasets, able to extract relevant representations, which can be repurposed for other task-specific predictions, such as peptides:MHC binding.

Here, we tested this hypothesis by building a CD4 T cell epitope predictor in which we modeled peptide:MHCII complexes with AF3 and processed the resulting structures using a graph-based neural network trained entirely on new in vitro binders identified by a highly-multiplexed assay.

## 2 Results

### 2.1 Generation of training data using a highly-multiplexed peptide:MHCII binding assay

To generate a new and diverse set of training data, we developed a system for highly-multiplexed in vitro peptide:MHCII binding assays. The PepSeq platform uses bulk in vitro transcription and translation to enable efficient preparation of programmable libraries of thousands to hundreds of thousands of DNA-barcoded peptides that can be used to perform highly-multiplexed assays [12]. We have previously described its application to high dimensional serology [16, 19]. Here, we describe MHCII-PepSeq, an adaptation of PepSeq that enables the analysis of peptide binding to MHC II proteins. In this assay, DNA-barcoded peptide libraries are incubated with purified MHC II proteins that contain a Class II-associated Invariant chain Peptide (CLIP) tethered by a cleavable linker [17, 37], in the presence of HLA-DM (Human Leukocyte Antigen DM isotype). After washing, peptides are quantified by amplification and sequencing of their DNA tags. Peptides that bind to the MHC groove are identified by a comparative analysis between two conditions: one in which the MHC is pre-treated with protease to cleave off the CLIP (‘cleaved’), and another in which MHC is left untreated, leaving the CLIP tethered (‘uncleaved’) (Figure 1 a).

**Figure 1.**
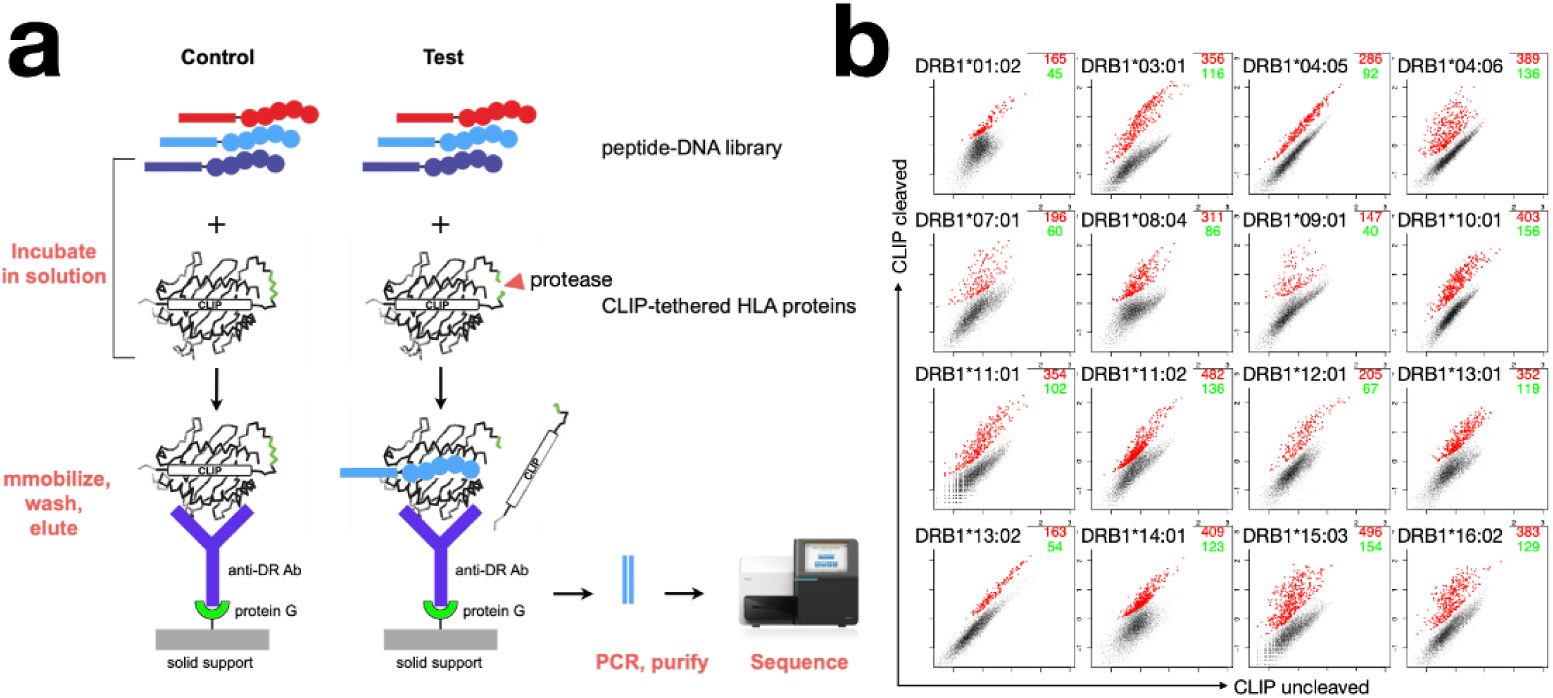
MHCII-PepSeq: a highly-multiplexed binding assay enabling epitope mapping across diverse MHC II proteins. **(a)** To enable highly parallel peptide:MHCII binding assays, libraries of DNA-barcoded peptides are prepared from DNA templates using bulk enzymatic reactions via the fully-in vitro PepSeq platform24. These libraries are then incubated in solution with recombinantly-expressed MHC proteins containing the groove-binding CLIP peptide tethered by a cleavable linker (either uncleaved – left, or cleaved with protease – right). MHCs and their bound DNA-barcoded peptides are then immobilized on magnetic beads, non-binders are washed away, and binders detected by amplification and sequencing of their DNA barcodes. **(b)** The MHCII-PepSeq assay described in (a) was used to assay a library of 9,003 overlapping 15mer peptides tiling 13 proteins from Plasmodium falciparum, across 16 HLA-DR proteins. Plots compare the sample-depth normalized abundances of peptides (1 dot per peptide) in the uncleaved (x-axis) and cleaved (y-axis) states. Peptides significantly enriched in the cleaved condition (by Fisher’s exact test) are highlighted in red. Numbers in the upper-right corners indicate the total number of enriched peptides (red), and the number of 9-13mer epitopes that are supported by ≥2 overlapping peptides (blue).

We generated a PepSeq library consisting of 9,003 unique 15mer peptides, tiling 13 Plasmodium falciparum proteins, in which each peptide overlaps its neighbor by 13 amino acids (step size = 2 amino acids). Using this library, we performed binding assays against 16 HLA-DR proteins. Comparison of ‘cleaved’ and ‘uncleaved’ conditions for each HLA using a Fisher’s exact test (see Methods) revealed distinct populations of 163-496 binding peptides per allele (total = 5,097) (Figure 1 b, red dots/values). Since the variable region of each PepSeq peptide is flanked by sequences of constant amino acids24 with the potential to contribute to MHC binding, to ensure stringency we focused our analysis on substrings of ≥ 9 amino acids that were each supported by ≥ 2 independent overlapping binding peptides (hereafter described as “epitopes”). By virtue of our peptide design parameters (15mers with step = 2), such epitopes were either 13, 11 or 9 amino acids in length. Using this approach, we identified 40-156 epitopes per HLA (total = 1,615) (Figure 1 b, blue values).

We compared these outputs of MHC-PepSeq with the predictions of a leading in silico peptide:MHC binding model, by generating Eluted Ligand (EL) scores using NetMHCIIpan4.3 [28] for the 1,615 PepSeq-identified epitopes across 16 HLAs described above. Eluted Ligand (EL) status corresponds to binary class that indicates whether a peptide is naturally processed by the MHC class II molecule. Score EL is a metric generated by NetMHCIIpan4.3 corresponding to the normalized predicted probability (via ranking) of being an elutable ligand. To enable fair comparisons, we ran these predictions on 15mer sequences centered on each epitope, and used a set of allele-specific negative 15mers (peptides that contained no 9mer in common with any binder peptide) as controls. Receiver operating characteristic (ROC) analysis treating PepSeq status as a binary variable and the NetMHCIIpan prediction as a quantitative metric revealed a range of concordances between the two methods across alleles, with AUCs from 0.75 – 0.97 (Figure 2 a). For 10 alleles, we observed AUCs ≥ 0.9. Overall, the degree of allele-specific concordance between PepSeq and NetMHCIIpan results appears to be (non-significantly, Pearson p=0.13) correlated with the size of the NetMHCIIpan training data for each allele (Figure 2 b). The most parsimonious interpretation of these findings is that PepSeq and NetMHCIIpan both accurately identify the same true binders for alleles that are well-studied, but PepSeq outperforms NetMHCIIpan on less well-studied alleles.

**Figure 2.**
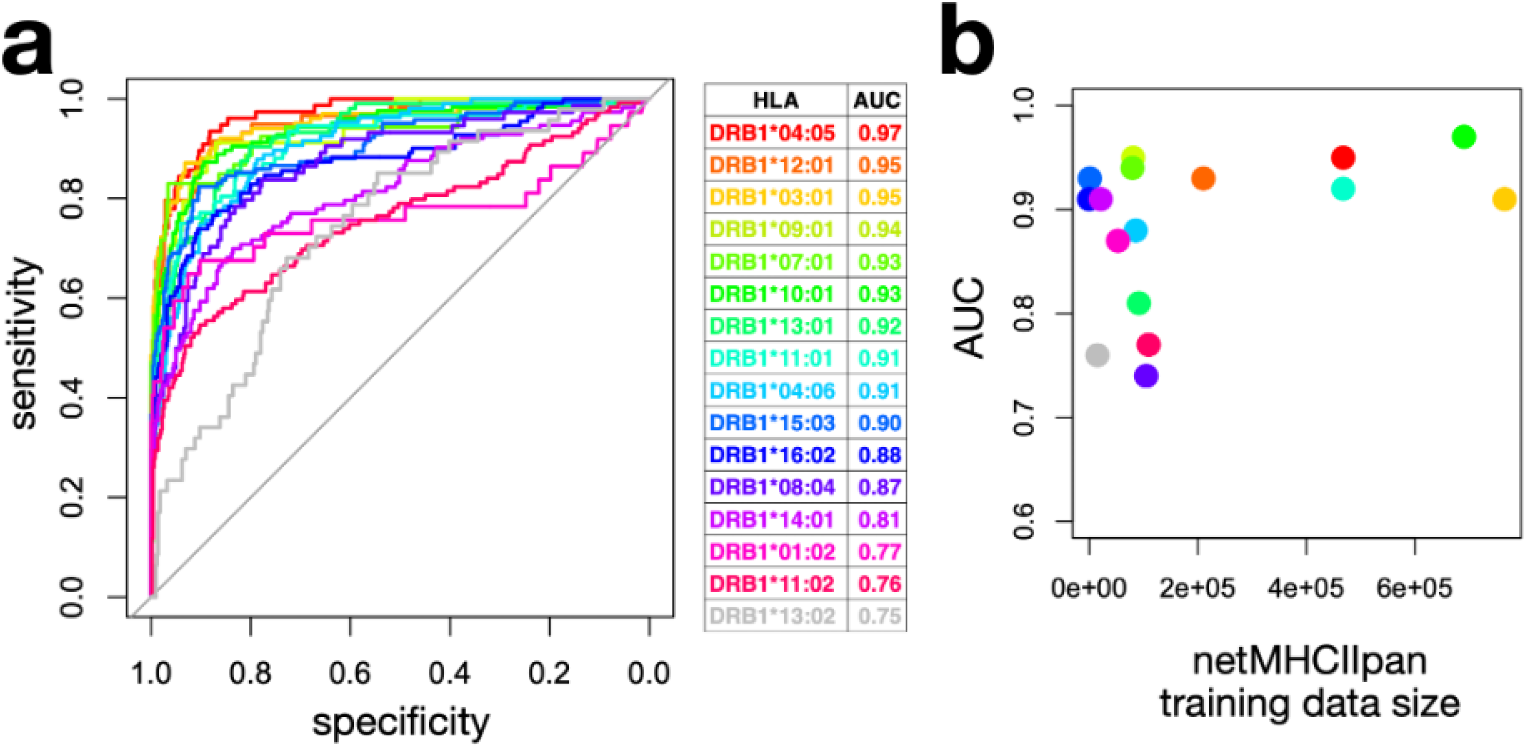
PepSeq and NetMHCIIpan are correlated, but diverge for a subset of under-trained alleles. **(a)** ROC analysis showing the degree of concordance between PepSeq and NetMHCIIpan4.3 EL for the assay data described in Figure 1b (9,003 peptides X 16 MHCs). NetMHCIIpan4 scores were calculated for peptide:MHC epitope pairs identified by PepSeq, as well as non-binding controls. **(b)** Correlation between size of the NetMHCIIpan training data and the AUC described in (a) for each of the 16 alleles. (a) Predicted structure of a peptide-HLA complex for a known weak binder. The peptide’s atoms are predominantly colored in bright yellow hues, indicating low pLDDT scores and reflecting the model’s low structural confidence in the peptide’s placement. (b) Predicted structure of a peptide-HLA complex for a known strong binder. The peptide is colored in dark blue, corresponding to high pLDDT values and strong structural confidence in the predicted interface.

Together, these results describe a new and scalable platform for high-dimensional MHC class II:peptide binding studies and its application to identify novel epitopes across diverse and understudied MHC proteins, useful for training new peptide:MHC binding models.

### 2.2 Classifiers for Peptide–HLA Binding Prediction Using AlphaFold3 Features

The predicted local distance difference test (pLDDT) is a confidence score between 0 and 100 for each atom in the predicted structure generated by AF3. It estimates how likely is the correctness of the predicted local atomic environment. The contact probability matrix is a square symmetric matrix where the entry (i,j) represents the contact probability between token i and token j, where a token (single amino acid in our case) is the basic unit of input sequence(s). By combining pLDDT-based metrics with the contact matrix-derived interaction features, we treat AlphaFold3 as a frozen encoder that transforms raw sequence input into biologically grounded structural representations. These representations serve as input to a downstream supervised model trained to predict peptide-HLA binding strength. In addition to the trained model, we also evaluate the predictive utility of the AlphaFold3-derived features in a zero-shot setting [24], where binding strength is estimated directly from the structural confidence metrics without any model training. Therefore, we evaluate two complementary classifiers for predicting peptide-HLA binding strength using AlphaFold3-derived structural representations.

The first is a zero-shot scoring approach that requires no additional training. For each peptide-HLA complex, we compute the value 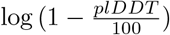 for every atom in the peptide and take the negative average over all peptide atoms as the final binding score. We will refer to this metric as the Fisher Confidence Metric (FCM) for its similarity with the Fisher’s method for combining p-values in a multiple testing setting [6]. The log-transformation amplifies the influence of atoms with high plDDT scores in the averaging process, based on our hypothesis that a peptide will form a strong binding interface if a significant subset of its atoms are confidently resolved in the predicted complex (Figure 3). This zero-shot score, that we refer to as the confidence metric, serves as an interpretable, confidence-weighted proxy for binding strength, derived entirely from the structure prediction model’s internal uncertainty.

**Figure 3.**
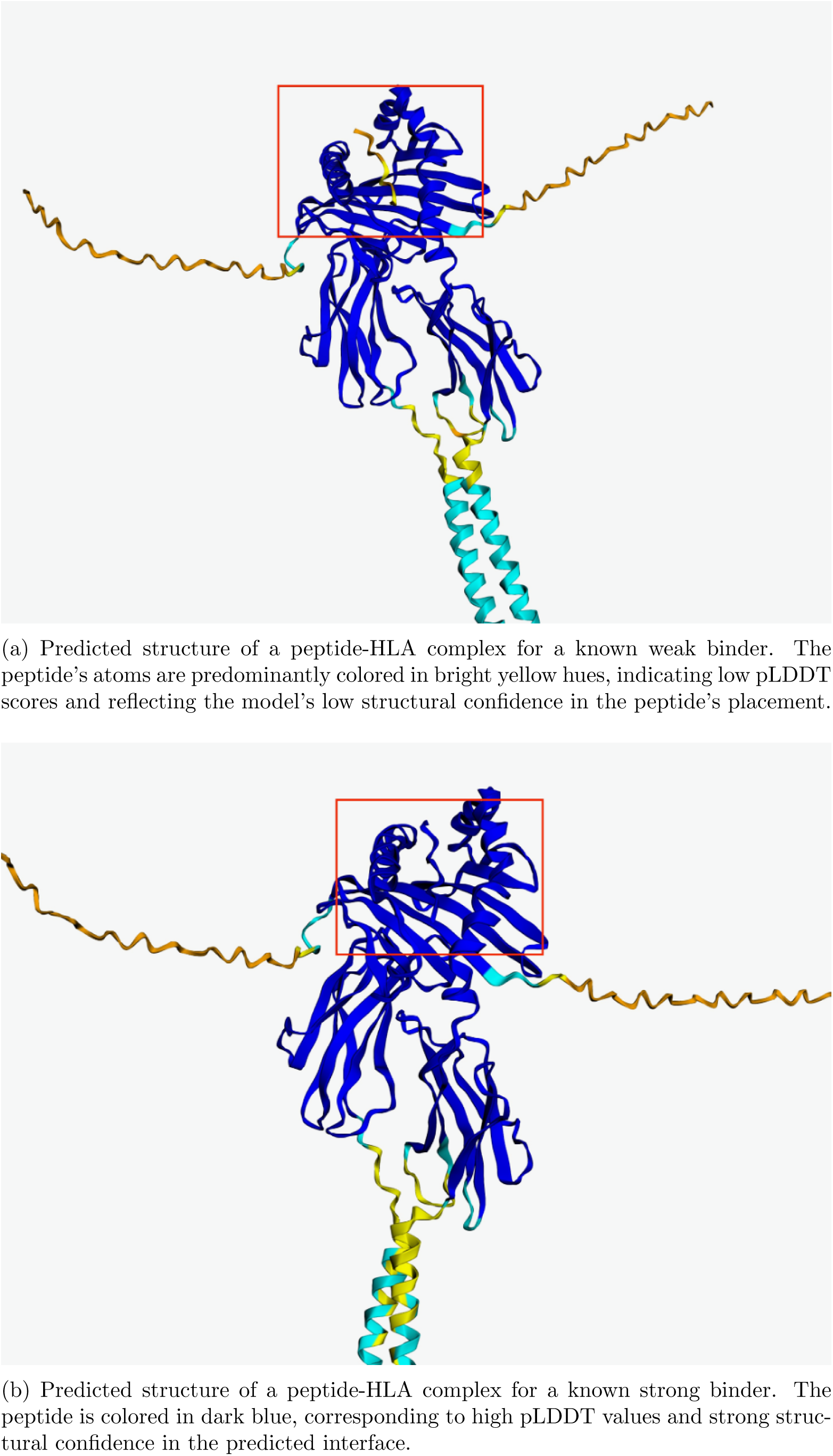
Visualization of AlphaFold3-predicted peptide-HLA structures for weak (a) and strong (b) binders. Color indicates pLDDT confidence scores per atom, with yellow representing low and blue representing high confidence. The structural confidence correlates with known binding affinities.

The second classifier adopts a transfer learning approach, using AlphaFold3 as a frozen encoder that supplies node-and edge-level features to a trainable Graph Neural Network (GNN). In this representation, each node corresponds to a token from the input complex—either a residue from the peptide or from one of the HLA chains. Node features include pLDDT scores and sequence/structure embeddings, while edges encode spatial proximity and contact probability. This framework enables the model to learn flexible patterns of interaction across the peptide-HLA interface while leveraging the pretrained structural knowledge embedded in AlphaFold3’s outputs. We will refer to our model as MHCIIFold-GNN. We describe the architecture and training procedure of the MHCIIFold-GNN classifier in detail in section 4.

Finally, we also combine the MHCIIFold-GNN with NetMHCIIpan 4.3 into an ensemble model and use it as a third classifier. To combine the GNNs-based scores with NetMHCIIpan binding scores, we used the geometric mean of the 3 scores involved: the predicted value from the amino acid type GNN, the predicted value from the confidence-based GNN and the NetMHCIIpan 4.3 Rank EL score (Rank EL/100)

### 2.3 Benchmark dataset

For both validation and testing, we use the benchmark dataset described recently for the evaluation of NetMHCIIpan 4.3 [22]. This dataset consists of CD4 T cell epitopes derived from the Immune Epitope Database (IEDB) [36], each associated with the restricting HLA protein, as well as the protein of origin of the epitope. Because there are only binders, an evaluation of these epitopes scores without a comparison to controls does not give any insight regarding a model performance. To evaluate the model’s ability to distinguish true MHC class II binders from background peptides, we adopt a ranking-based evaluation approach corresponding to the FRANK value metric described previously by [22]. For each test epitope, we retrieve its source protein and generate a set of control peptides by sliding a window of the same length as the epitope across the entire protein sequence, and generate corresponding scores for each control and the tested epitope. Specifically, if the epitope is of length L and the source protein is of length N, we generate N-L overlapping peptides of length L, excluding the exact match to the epitope itself (Figure 4). Each of these peptides is assigned a predicted binding score using our model, and the epitope is then ranked relative to its corresponding background set. This procedure provides a protein-specific contextual evaluation, assessing whether the model assigns higher binding scores to the true epitope compared to all other same-length peptides from the same source protein. The FRANK value corresponds to the fraction of control peptides with a higher predicted binding score than the true epitope, and serves as a summary statistic of model performance for every positive epitope.

**Figure 4.**
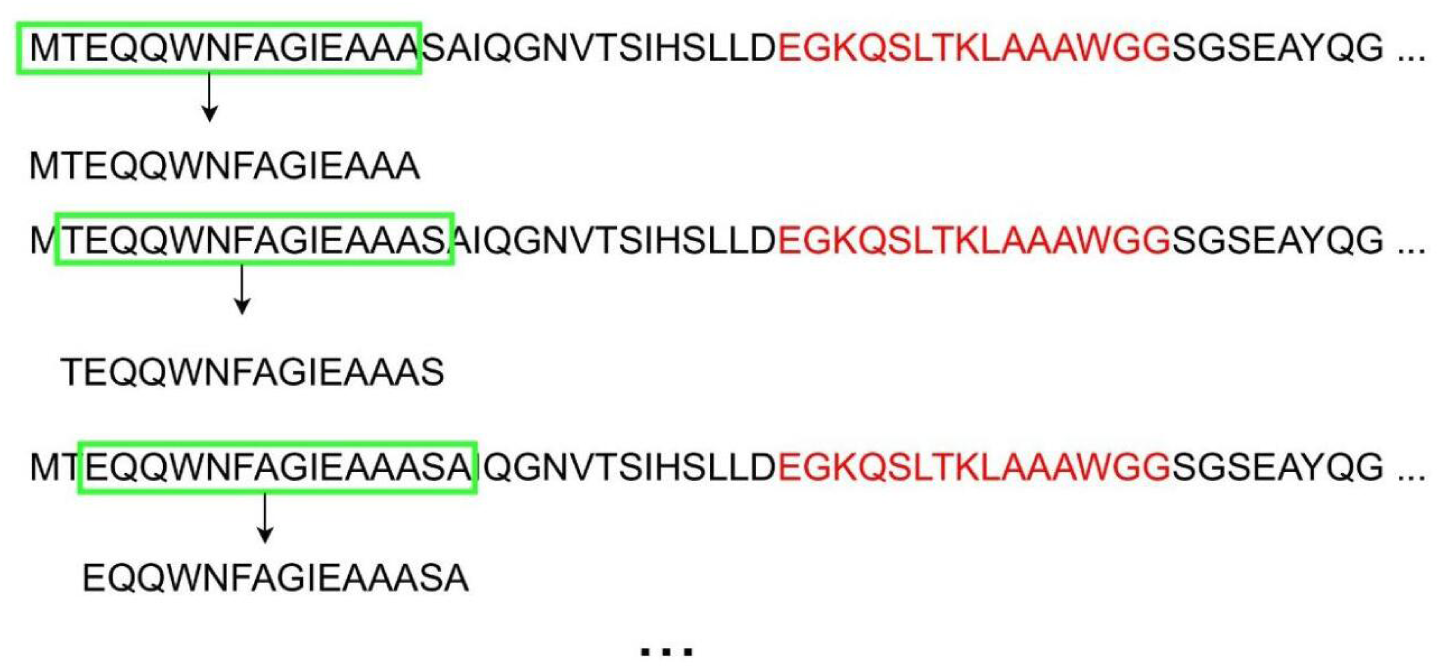
Background peptides generation: sliding windows of the same length as the epitope (in red) across the entire protein sequence are generated, excluding the red window.

For our validation dataset and test sets, we focused exclusively on alleles of the HLA-DRB1 locus, corresponding to the HLA gene studied by PepSeq. We randomly selected 23 out of the 34 DRB1 HLA alleles available in the dataset for analysis, and 7 HLAs of the 23 HLAs to form our validation set.

To compare our results to NetMHCIIpan 4.3 results, we ran the same epitopes and negatives used to evaluate the performance of NetMHCIIpan4.3. Using the Rank EL metric, we computed FRANK values, enabling one-to-one comparisons between our results and NetMHCIIpan results.

### 2.4 Models Performance Comparison on Benchmark Dataset

To assess the effectiveness of AF3-derived structural features for CD4 T cell epitope prediction, we evaluated the four models on a held-out test set:

i. the established NetMHCIIpan 4.3 model
ii. the zero-shot confidence metric
iii. the GNNs-based model where we input frozen features from AF3 and train the GNNs via labeled PepSeq epitopes/nonbinders
iv. the combination of the MHCIIFold-GNN and NetMHCIIpan 4.3.

In Figure 5, we present the mean FRANK value observed for each of the four methods under consideration.

**Figure 5.**
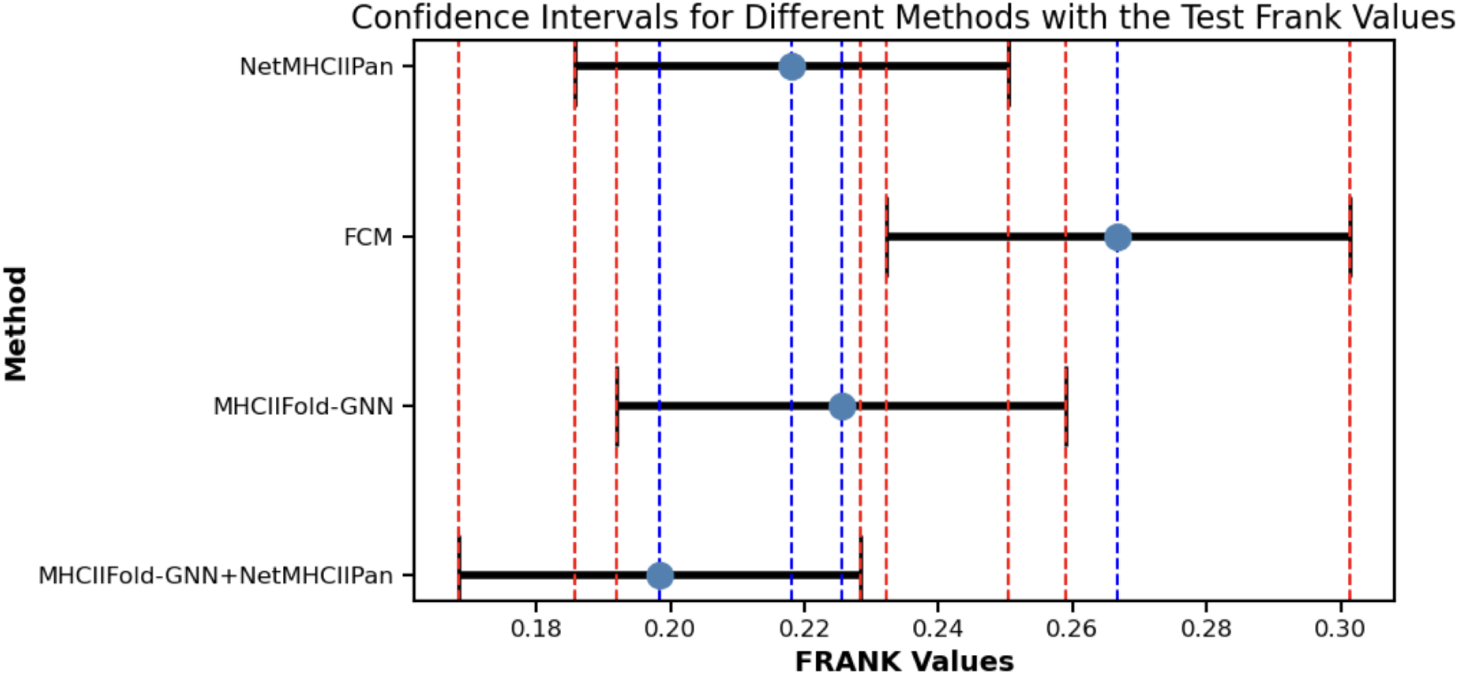
Confidence intervals and the means of all the Frank Values of each method

Across all epitopes, NetMHCIIpan yielded a mean FRANK value of 0.218 (95 % CI [0.186, 0.250]). Despite its simplicity, the zero-shot confidence metric, which estimates binding directly from pLDDT scores without any supervised training, achieved a mean FRANK value of 0.267 (95 % CI [0.232, 0.30]). This model already performs competitively with NetMHCpan 4.3. This result underscores the relevance of AF3 structural confidence output alone in the peptide:MHC binding problem.

To assess the added value of a trainable architecture on top of these structural representations, using a transfer learning approach, we evaluated our GNNs-based model. The MHCIIFold-GNN achieved a lower mean FRANK value of 0.225 (95 % CI [0.191, 0.259]), demonstrating that fine-tuning on a small labeled dataset further improves performance over the zero-shot baseline. Finally, the combination of the MHCIIFold-GNN and NetMHCIIpan 4.3 yields a further improvement, achieving a mean FRANK value of 0.198 (95 % CI: [0.168, 0.228]). In other words, integrating AF3-derived structural features with existing prediction frameworks via an ensemble model achieves more than 11 % improvement over NetMHCIIpan 4.3 alone.

To further illustrate the FRANK scores distributional characteristics of each method, Figure 6 shows boxplots of the FRANK values across all epitopes, enabling comparison of performance variability and robustness. In Figure 7, these boxplots are stratified by HLA allele, revealing how predictive performance differs across specific MHC class II molecules.

**Figure 6.**
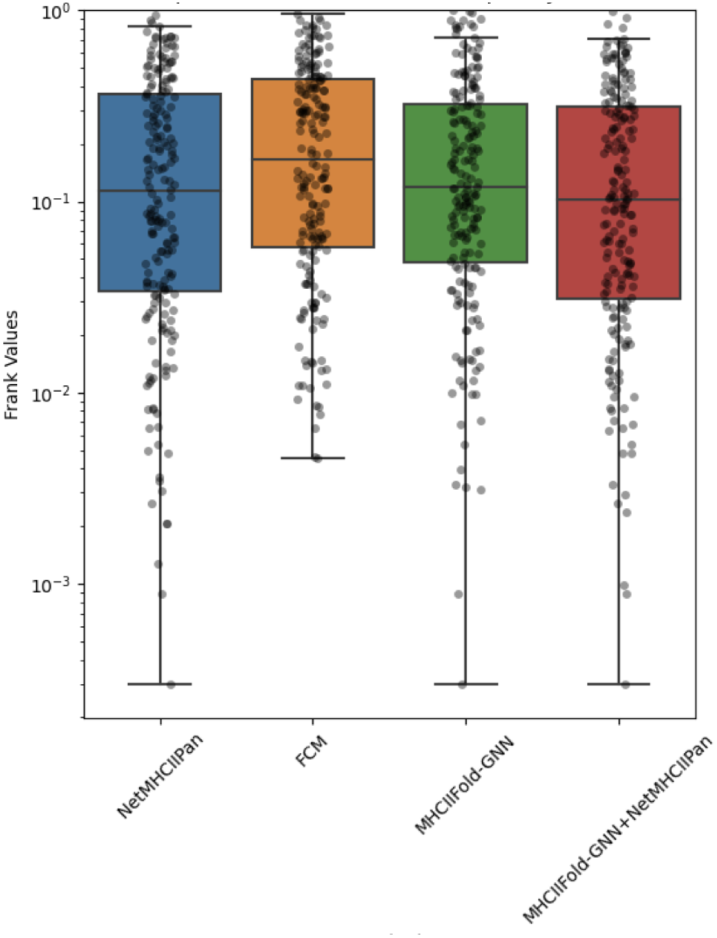
Distribution of FRANK values across all epitopes for each method.

**Figure 7.**
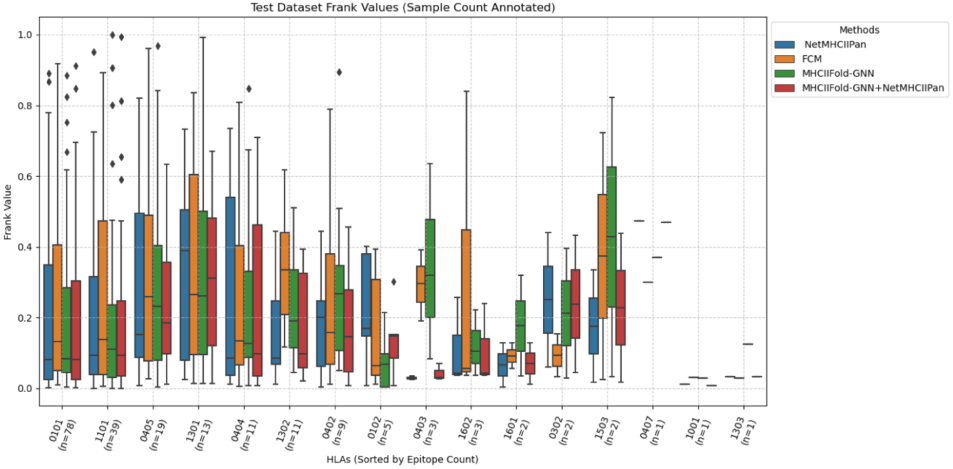
Distribution of FRANK values stratified by HLA allele.

When epitopes FRANK values are compared pairwise via a two tailed paired Wilcoxon test, we observe a significant p-value between the NetMHCIIPan 4.3 results and the combined model results (p = 0.0005), highlighting the added value of our model when combined with NetMHCIIPan. However, the test did not yield a significant p-value between the MHCIIFold-GNN and the combined model (p = 0.38).

## 3 Discussion

Our findings establish that structure-based modeling using AlphaFold3-derived features, combined with a lightweight graph neural network, enables accurate prediction of peptide:MHC class II binding across diverse alleles, even in the absence of ligand elution training sets. Remarkably, our GNN-based model, trained exclusively on in vitro binders from a multiplexed assay, performs competitively with NetMHCpan 4.3-a leading machine learning model trained on vast and heterogeneous datasets. Compared to our PepSeq training set, which included ≈52K non-binders and 633 binders, NetMHCIIpan required over 7 million peptides—more than 100 times larger—to achieve comparable performance (See section 4.3). This result underscores the expressive power of AF3 structural representations for immunological applications and highlights peptide:MHC binding as a central determinant of CD4 T cell epitope recognition.

In addition to its utility in the training of MHCIIFold-GNN, our development here of MHCII-PepSeq as a platform for assaying thousands of fully-customizable peptides for binding across diverse MHC class II proteins promises to have a number of future applications. First, in cases where maximum accuracy is important and/or time and cost are less limiting, custom PepSeq libraries can be generated to query particular peptide sets of interest for MHC binding experimentally. Second, we expect the analysis of larger PepSeq libraries, and more diverse MHC alleles (including many that remain vastly understudied), in combination with learning approaches such as the structural one described here, will lead to further improvements in the accuracy of predictive MHC binding models. While it does not address the upstream steps involved in antigen processing, MHCII-PepSeq offers several attractive features compared to ligand elution assays, including unambiguous assignment of MHC alleles, and experimental simplicity and scalability. Relatedly, we expect PepSeq analysis of MHCII binding to scale beyond the 10,000 peptide library demonstrated here, likely tointo the ≥ 100000s of peptides rangescale that we have demonstrated in the serological context [12, 19]. Notably, our focus on non-random peptide sequences distinguishes our approach from alternatives [11, 26], and may in the future enable an iterative learning cycle in which custom peptide sets are designed on the basis of prior predictive models, in a way that maximizes the rate of new learning.

A particularly noteworthy outcome is the performance of the zero-shot confidence metric. Despite requiring no training (other than training the AF3 model), this simple proxy–derived from pLDDT scores– achieves results comparable to those of NetMHCIIpan. This observation reinforces the biological value of AF3’s internal uncertainty estimates and offers a practical method for peptide prioritization when labeled training data is limited or unavailable. By applying a GNN-based model to AF3-derived structural outputs, we were able to further improve predictive performance, demonstrating the benefits of minimal fine-tuning in a transfer learning context. Importantly, our GNN captures a subset of epitopes that NetMHCIIpan scores as weak binders and vice versa. This complementarity is reflected in the improved performance of the ensemble model, which combines outputs from both frameworks and yields the best FRANK scores overall. These results suggest that AF3-based learning encodes orthogonal information that is not captured by existing sequence-based models, and can be effectively leveraged through appropriate graph-based representations.

Several limitations remain and there is clearly a need for further investigation. The number of unique peptides in our training set is relatively small, and the length distribution of training peptides is shorter than that of peptides in the IEDB epitopes dataset (Figure 8). This mismatch limits the number of windows contributing to each peptide and may hinderpenalize model generalization. Moreover, the diversity of HLA alleles included in training was restricted to DRB1 types for which high-quality PepSeq data was available. As a result, our method was only tested on DRB1 alleles as well. Training and testing in other alleles remains to be done to support the robustness and transferability of structural AF3 features.

**Figure 8.**
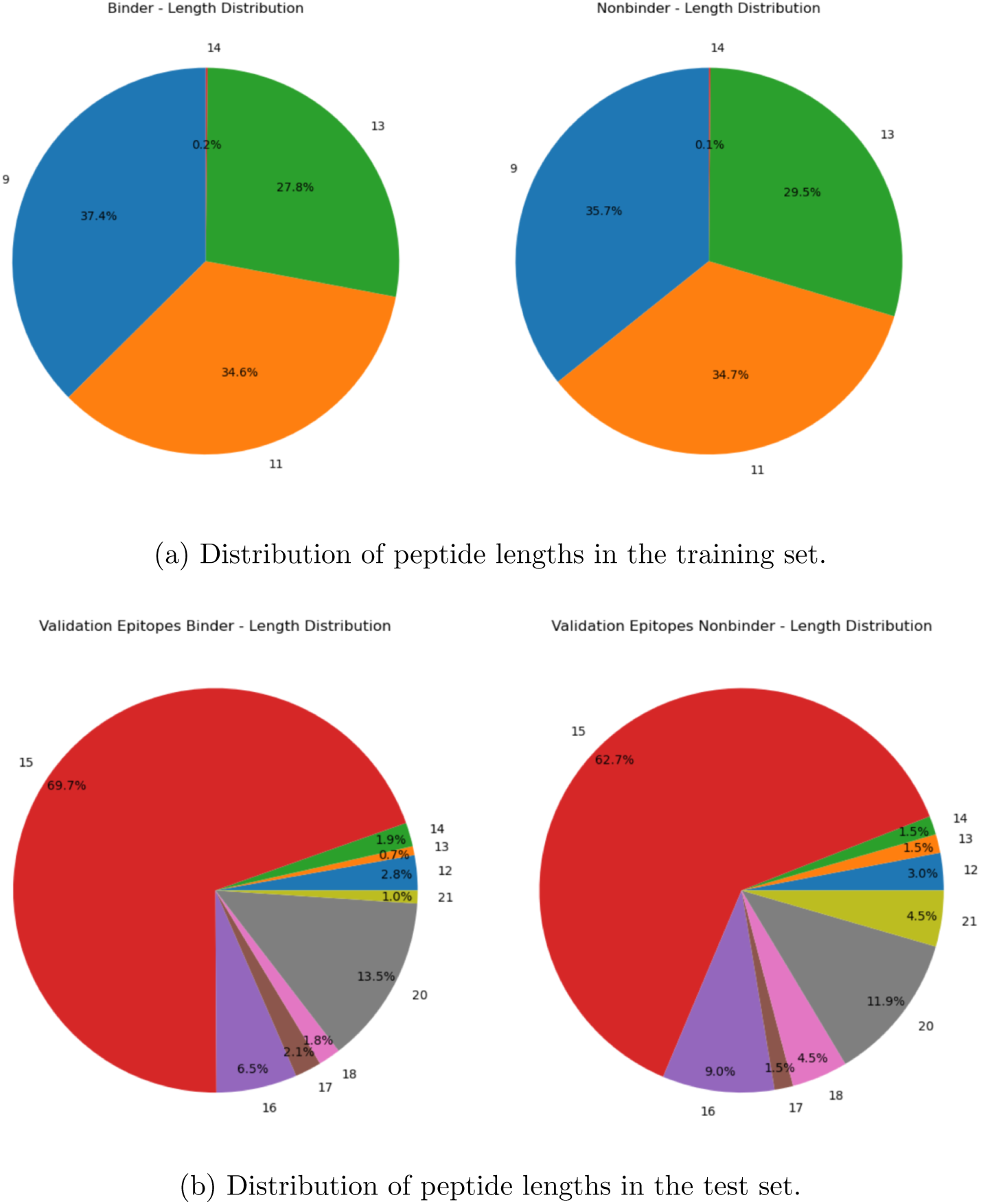
Distributions of peptide lengths in the dataset. (a) Training set. (b) Test set.

Importantly, while the combined models show improved mean FRANK scores, the 95 % confidence intervals for all evaluated methods overlap, indicating that no model is statistically significantly better than the others given the current test set (number of tested epitopes) and its size. This underscores the need for larger and more diverse test sets to achieve greater statistical power and draw more definitive conclusions regarding comparative performance.

Looking ahead, we anticipate that model performance could improve further with expanded training datasets–including both larger sets of characterized sequences and broader HLA coverage– and with integration of naturally processed ligands from immunopeptidomics studies. Future work may also explore whether combining structure-informed learning with experimentally validated T cell epitope data could support more holistic models of antigenicity, extending beyond binding. In parallel, designing deeper and more expressive GNN architectures trained on larger and more HLA-diverse datasets may further enhance prediction accuracy, improve generalization to rare alleles, and expand applicability to clinical settings.

More broadly, this study provides a conceptual justification for structure-informed prediction pipelines using large foundation models like AF3. By treating AF3 as a frozen encoder, we make the case for its intermediate structural representations to be repurposed effectively for downstream prediction tasks, enabling high performance even in data-limited settings. In future work, fine-tuning intermediate layers of AF3 – rather than using it purely as a feature extractor – may enable the model to learn binding-relevant structural adjustments specific to the peptide:MHC interface.

These directions point to an emerging paradigm: Using pre-trained powerful foundation models such as AF3 with simpler task-specific neural architectures to advance complex fields such as precision immunology.

## 4 Methods

### 4.1 PepSeq library design and synthesis

We designed a library of 9,003 15mer peptides tiling the sequences of 13 Plasmodium falciparum proteins (NCBI protein accessions = CAD51981, CZT99892, CZT99492, CZT99491, CAD49145, CZT99074, XP 001351122, CAD52497, CAD52483, CAD52255, CZT98878, CAG25193, CAD48982, of which 11 were selected for preferential expression during the liver stage of infection). Tiles were constructed with a step of 2 amino acids, such that each peptide overlaps its neighbor by 13 amino acids. The designed set of peptides was encoded into DNA and used as templates for the production of DNA-barcoded peptide (PepSeq) libraries, as previously described [12]. Briefly, from 48 pM of the commercially synthesized peptide-encoding ssDNA oligonucleotide library (Agilent Technologies, USA), a PCR was performed using a specific primer pair to amplify dsDNA sequences. The resulting PCR products were then utilized to synthesize strands of mRNA in an in vitro transcription reaction using the AmpliScribe T7-Flash system (LGC Biosearch Technologies, USA). The mRNA products were ligated to a hairpin oligonucleotide adaptor that contained a puromycin molecule linked by a polyethylene glycol (PEG) spacer.

Then, the resulting mRNA-puromycin complex served as a template in a cell-free translation reaction employing the PURExpress protein synthesis system (New England Biolabs, USA). Afterward, a reverse transcription reaction primed by the adaptor hairpin was utilized to produce cDNA. The original mRNA was degraded using RNase H (New England Biolabs, USA) and an RNase cocktail (Invitrogen, USA), leaving the synthesized peptide attached to the cDNA. Finally, the resulting cDNA-barcoded peptide library was treated with a tobacco etch virus (TEV) protease (Invitrogen, USA) to remove the constant region amino acids from the N-terminal end of the peptides.

### 4.2 MHCII-PepSeq binding assay

HLAs were sourced from the Tetramer Core Facility (under contract 75N93020D00005) and incubated overnight at room temperature with either thrombin or 3C protease (Millipore Sigma, USA) to release the tethered CLIP peptide. Next, 1 pmol of the PepSeq library was mixed and incubated overnight at 30°C with 0.5*µg* of each cleaved or uncleaved (negative control) HLA and 2*µg* of HLA-DM. The binding buffer contained 230 mM NaCl, 110 mM citrate buffer pH 5.5, 4.7 mM EDTA pH 8.0, 0.9% n-Octylglucoside, and 1.3% protease inhibitor cocktail (Promega, USA). These binding reactions were done in triplicate. After incubation, HLAs were captured using an anti-human HLA DRB1-L243 antibody (BioLegend, USA) bound to protein G-bearing beads (Invitrogen, USA) for one hour at room temperature. Samples were washed, eluted, and retrieved from the beads. Finally, samples were amplified and indexed by PCR as previously described (Henson et al., 2023). After PCR cleanup, the products were pooled equally and quantified. Finally, 800 pM of the pooled library was loaded and sequenced using an Illumina NextSeq 1000 instrument.

### 4.3 MHCII-PepSeq data analysis

We aligned sequencing reads to the library of DNA templates using the PepSIRF software package described previously [8], using the demux module with default parameters. To identify binding peptide for each HLA allele, we next applied a Fisher’s test method in which, for each focal peptide in the library, we tested a contingency table of mapped read counts for the ‘cleaved’ v ‘uncleaved’ conditions for the focal peptide v the sum of all other peptides. Allele-specific p-value thresholds were selected that best captured the visual separation between populations (per Figure 1 b), and these values were fixed prior to all downstream analyses. Next, we applied an “intersection rule” to identify substrings of ≥9 amino acids that were each supported by ≥2 independent overlapping binding peptides. We also identified allele-specific negative 15mers (peptides that contained no 9mer in common with any binder peptide) as controls for the ROC analysis. We focused GNN training on the top HLAs with the highest concordance with NetMHCIIpan but with different numbers of HLAs for the different parts of the MHCIIFold-GNN model (Figure 2 b).

After choosing the PepSeq epitopes and nonbinders based on the intersection rule explained above, the data needed one more processing step to remove length biases. Because of the intersection processing, the PepSeq epitopes had lengths between 9-13, but all the PepSeq nonbinders had a length of 15 amino acids. This would create a problem for the model. To make the length distribution comparablereasonable between PepSeq epitopes and nonbinders, we used the following assumption: because PepSeq nonbinders are relatively short, removing a short portion from these peptides won’t affect their binding positively, i.e. they would still be PepSeq nonbinders. Following this assumption, we trimmed the 15mer PepSeq nonbinders from one of the ends. The PepSeq nonbinders, the end they will be trimmed from, the length of the trim were all random. This processing would equalize the length distribution between the classes. So, if the length distribution of PepSeq epitopes in an HLA was 1:3:1 (1 for 9mers, 3 for 11mers, 1 for 13mers), the PepSeq nonbinders would have the 1:3:1 distribution as well (Figure 8a). After trimming peptides to match the length distributions between PepSeq epitopes and nonbinders as described, some PepSeq nonbinder sequences become identical (duplicates). The duplicate peptides are removed for every given HLA.

Our training set had 633 PepSeq epitopes, 51534 PepSeq nonbinders, yielding 150131 total 9mer graphs and 112530 total unique 9mer graphs across all HLAs combined.

As a comparison, the EL NetMHCIIpan training set is had total of 675,364 positive and 6,886,973 negative peptides from a total of 237 EL samples, covering a total of 142 MHC class II molecules. Furthermore, the binding affinity (BA) data consists of 129,110 data points covering 80 class II molecules. An overview of all the datasets used in the study in terms of peptide counts, HLA types, dataset type (BA and EL) and processing method (preprocessed or filtered) are provided in table S1.

### 4.4 AF3-derived structural features for model construction Structural Input Configuration

Each input to AlphaFold3 comprises the two chains of the HLA heterodimer (*α* and *β* chains) and a candidate peptide typically of length 8-20 amino acids. AlphaFold3 is then run in complex prediction mode to generate the 3D structure of the full peptide-HLA assembly together with auxiliary features. We next describe the relevant auxiliary features generated by AF3 that were used in our procedure.

#### Predicted local distance difference test (pLDDT)

pLDDT is a confidence score between 0 and 100 for each atom in the predicted structure. It estimates how likely is the correctness of the predicted local atomic environment. For example pLDDT values ≥ 90 indicate a very high confidence that the corresponding atom is placed correctly [1]. The idea is to use the pLDDT score as a proxy for structural stability. Indeed, we hypothesize that high pLDDT scores across a contiguous substring of the peptide indicate more stable interactions, hence the interest in using these scores as individual per atom features, and aggregate them using a downstream model for our peptide-HLA binding prediction task.

#### Contact Probability Matrix

The contact probability matrix is a square symmetric matrix where the entry (*i, j*) represents the contact probability between token *i* and token *j*, where a token is the basic unit of each input sequence [1]. A token is simply an Amino Acid in our case, where the input sequences are proteins. A contact between tokens is defined as a distance less than 8Å between the representative atoms of a token. In our case, the matrix captures the model’s predicted likelihood of physical proximity between each pair of residues in the complex. We extract the submatrix corresponding to interactions between the peptide and the HLA chains, which reflects the model’s inferred binding interface. The matrix can serve as a geometry-informed descriptor of the interaction landscape, encoding the presence and significance across the peptide-HLA complex.

### 4.5 MHCIIFold-GNN

We propose a model based on two graph neural networks (GNNs) [38, 32] to predict the binding strength of the MHC II proteins and peptide chains. For reasons that will become clear when describing the two GNNs below, we will refer to the first GNN as the confidence-based GNN and the second as the amino acid-type GNN. Each GNN model outputs one predicted score and these scores are combined via their geometric mean, yielding one score. Each peptide is broken into 9mer windows (9mer reference) with a sliding window. The combined two GNN scores represent a binding score of a single 9mer window of a given peptide. For example, if a peptide has length 11, it would have 3 windows (Figure 9). Each of these 9mers is run separately through AF3 with its corresponding HLA alpha and beta chains. Each of the 3 windows is then processed through the two GNN models, one window at a time, to output one score per window. We obtain a peptide score per GNN after taking the geometric mean of all the peptides (9mer) windows scores (Figure 4a). We again combine each of the two GNN scores via their geometric mean to get the final peptide score (Figure 4b) The idea of combining the windows scores is related to Fisher’s method for combining p-values and has been used previously in NetMHCIIpan. We next present how the two GNNs model is constructed. The pseudo code detailing the full GNNs based training and inference is summarized in pseudo code in Appendix B.

**Figure 9.**
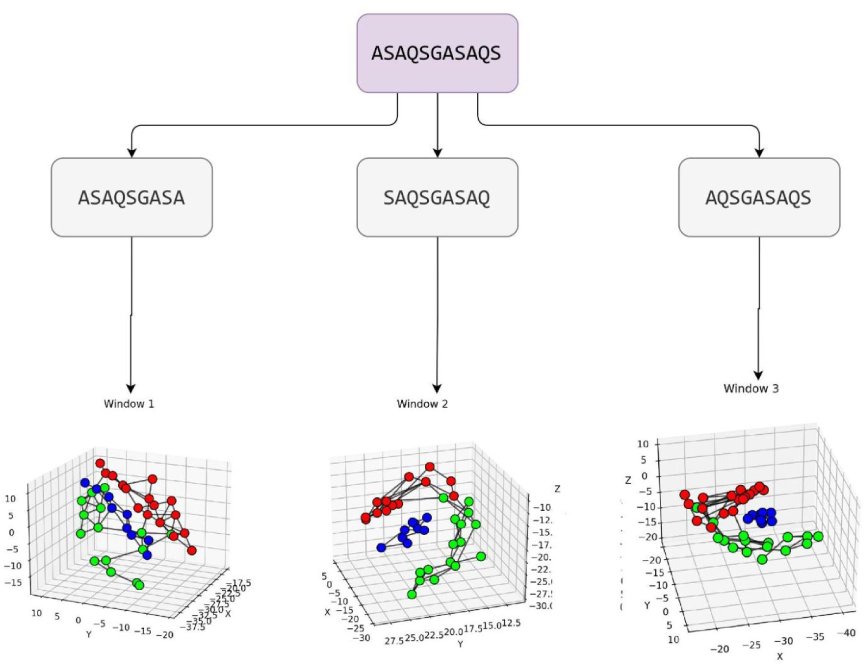
Illustration of the sliding window and model processing framework for an 11-mer peptide. The peptide is split into overlapping 9-mer windows (3 in this case), each paired with its corresponding HLA alpha and beta chains and processed independently through AlphaFold3 to obtain structural information. For each window, a graph is constructed and passed through the same two GNN models—(1) the confidence-based GNN and (2) the amino acid-type GNN—producing one binding score per window. These scores are aggregated via geometric mean to form peptide-level scores for each GNN, which are then again combined geometrically to yield the final binding score.

**Figure 10.**
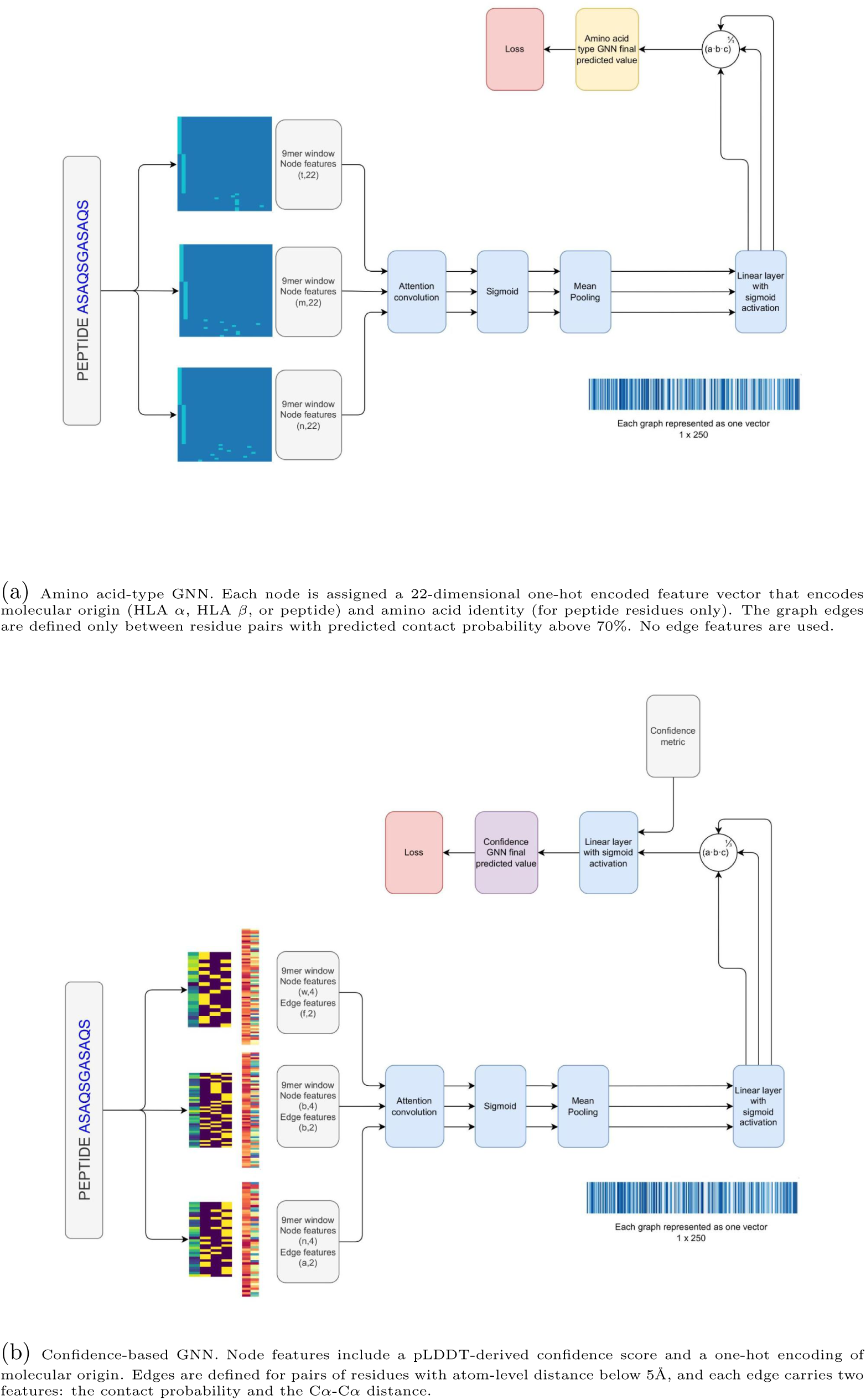
Input representations and architectural differences between the two GNNs used for peptide-HLA binding prediction. While both GNNs share the same overall structure and processing pipeline, they differ in how they define graph topology and represent input features.

#### Generating the input graphs of the GNNs

The two GNNs share a similar input graph structure. In both cases, the nodes of the graph represent amino acids coming from the peptide window or the HLA chains. For the HLA chain amino acids, only amino acids sharing an edge with a peptide amino acid are kept. We therefore HLA amino acids that are in close proximity to the peptide chain and play a potential role in the binding of the peptide. We use 3D coordinates of the atoms generated by AF3, and the contact probability matrix to generate edges between amino acids. For the confidence-based GNN, two amino acids have an edge if and only if they have a pair of residue atoms with a distance less than 5Ä. The input graph of the amino acid type GNN is the same, with the additional constraint of drawing an edge between two amino acids only if the AF3 predicted contact probability between the amino acids is greater than 70%.

#### Defining the input node and edge features of the GNNs

The confidence-based GNN has a nodes feature vector represented by 4 entries. The first entry is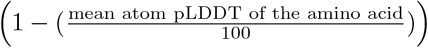 for the corresponding amino acid. The other 3 entries represent a one-hot encoded vector of the molecular type of the nodes: (1 0 0) for the HLA alpha chain amino acid, (0 1 0) for the HLA beta chain amino acid, and (0 0 1) for a peptide amino acid. The edge features have 2 entries: the contact probability for that edge and the distance between the alpha-carbons of the two corresponding amino acids. Since we create edges based on the residue atoms’ distances, it is possible some edges will have distance features larger than the 5Ä threshold.

For the amino acid type GNN, we only have node features. To encode molecular identity and amino acid type within our graph representation, we introduce a 22-dimensional one-hot node feature vector.. The first two entries indicate whether the amino acid belongs to the HLA *α*-chain or HLA *β*-chain, respectively. The remaining 20 entries represent a standard one-hot encoding of amino acid type, which is included only for residues originating from the peptide. For nodes corresponding to HLA *α* or *β* chains, all 20 amino acid-type entries are set to zero. This design acts as a form of structural regularization, allowing the model to rely on amino acid identity only in the peptide, where binding specificity is expected to be most critical, while encouraging the model to generalize across diverse HLA sequences without overfitting to specific residues in the *α* or *β* chains as we don’t have enough diversity of HLA chains in our training set.

#### GNNs architecture

Each of the 9-mer windows is in the proper structure for the GNN. However, before inputting them into the GNN, we combine the 9mers that belong to the same peptide in one group for both models. This way, the GNN will input the peptide, which will have multiple graphs but only one label for the loss computation.

The GNN has two components: one that works on individual 9mer windows and one that combines the values of the 9mers that belong to the same peptide.

Both GNNs are rather shallow, highlighting the relatively small number of layers needed to process the features extracted from AF3. In both GNNs, we start by applying one attention convolutional layer to the input graph with its node features and edge features. The next layer yields 250 dimensions via 10 multi-attention heads (25 dimensions per each head) using a graph convolutional layer with attention (GATv2Conv) [35, 3], followed by a sigmoid function and global mean pooling for both models. Finally, a linear layer with a sigmoid activation function is implemented to transform the 250-length vector into a scalar value. At this stage, we obtain a scalar value for each window. All window scores are combined by computing their geometric mean. The geometric mean gives more weight to 9mer windows that are string binders. For the confidence based GNN model, we have a final layer where the confidence metric and the output of the geometric mean layer are combined via a linear layer followed by a sigmoid activation. Finally, the two GNN scalar outputs are again combined via their geometric mean, yielding the final predicted score.

#### GNNs training

The two GNNs are trained separately. In each model, the cost function is binary cross-entropy loss where we have one unique label per peptide and the labels are binders/nonbinders. These labels are obtained from the PepSeqs runs. We used a standard inverse standard inverse class frequency weighting to balance the overrepresentation of nonbinders in the training set. We optimized the model using Adam optimizer. 14 HLAs were used for the amino acid based GNN while only 10 HLAs were used for the confidence based GNN. This choice was only made based the validation dataset and then applied to the test set. The number of HLAs for each model and number of epochs and each model was picked based on the mean of the confidence interval of the FRANK values of the validation dataset. The combination yielding the minimum mean value (after 30 epochs for all models) was chosen. Finally, 120 epochs were used for the training of the amino acid based GNN and 880 epochs were used for the confidence based GNN. The training dataset of the amino acid GNN has 1230 PepSeq epitopes, 97412 PepSeq nonbinders were used. For the confidence-based GNN, 895 PepSeq epitopes, 69825 PepSeq nonbinders were used. A summary of the HLA alleles used in training, validation, and testing, including the number of binders and nonbinders from the PepSeq training set and the epitope benchmark dataset, is provided in appendix A, Table 1.

#### Software environment and reproducibility

All analyses were conducted within a containerized environment based on Python 3.10. The container included the following key packages and versions: pandas 2.2.2, numpy 1.26.0, scipy 1.12.0, and scikit-learn 1.5.2. For graph neural network modeling, we used PyTorch 2.4.1, torchvision 0.19.1, torchaudio 2.4.1, and the PyTorch Geometric library (torch geometric 2.6.1), along with its dependencies: pyg-lib 0.4.0, torch-scatter 2.1.2, torch-sparse 0.6.18, torch-cluster 1.6.3, and torch-spline-conv 1.2.2. Additional packages used for analysis and data processing included pyarrow 18.0.0 and biopython 1.81. For visualization in Jupyter notebook, we used seaborn 0.12.2, matplotlib 3.7.1. Model training was performed using the Adam optimizer with a learning rate of 0.001. The confidence-based GNN was trained for 420 epochs with a batch size of 128, while the letter-model GNN was trained for 120 epochs with the same batch size. To ensure reproducibility, a fixed random seed was used across all relevant libraries, including PyTorch, NumPy. For structure inference and auxiliary features generation, AlphaFold version 3.0.1 (https://github.com/google-deepmind/alphafold3) was used. All (total 74,063 unique) 9mer peptide windows were run in –no run inference mode to generate multiple sequence alignments (MSAs), which were then merged with HLA alpha and beta chain MSAs and executed in – no run datapipeline mode with a fixed seed to generate the final structural predictions and auxiliary features. Inference jobs were managed using custom Python scripts and a Nextflow pipeline.

## A HLA/Peptides dataset table

**Table 1.**
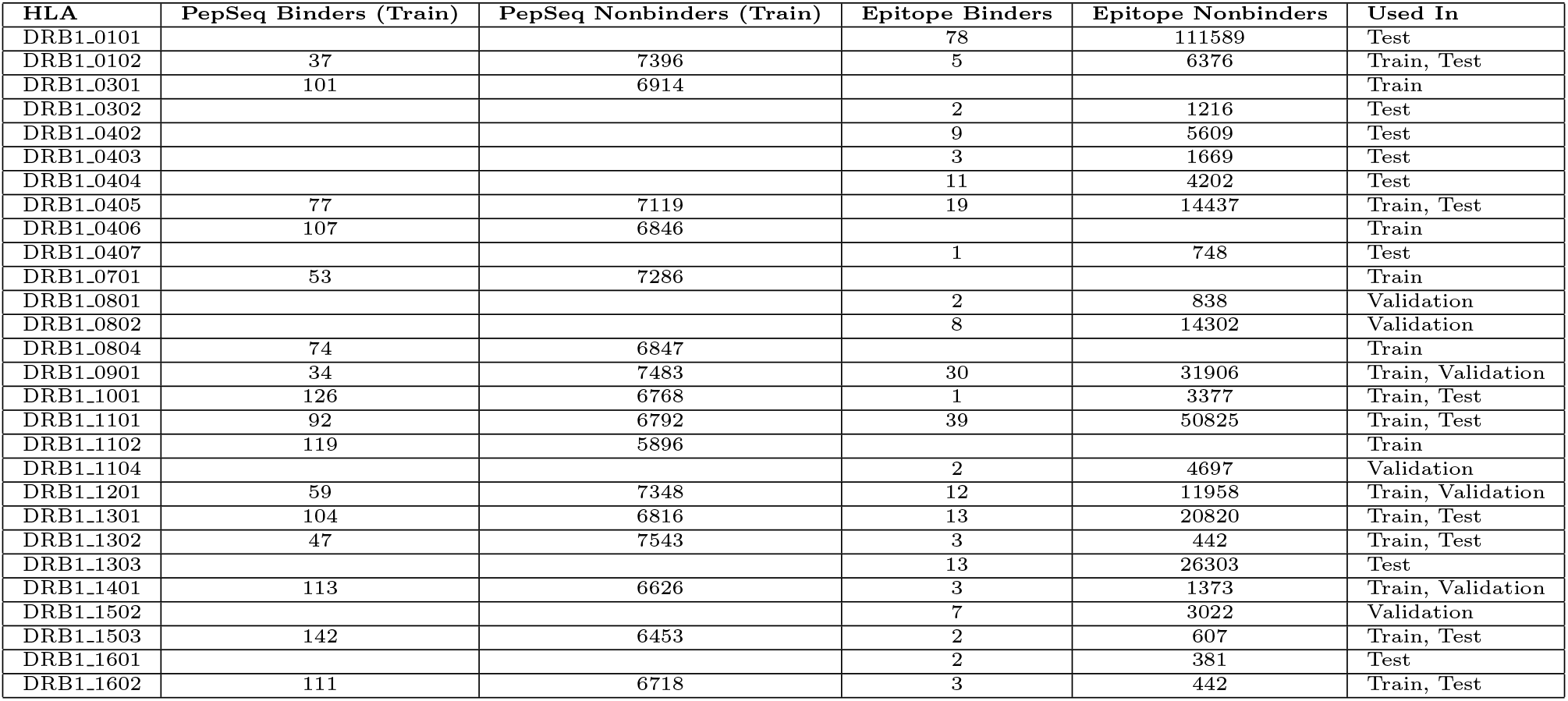
Summary of peptide counts for each HLA allele across training, validation, and test datasets. Alleles with PepSeq-derived binders or nonbinders were included in training. Alleles with epitope data were used for validation or testing, as indicated in the “Used In” column.

## B Pseudo code for MHCIIFoldNN

### Algorithm 1

MHCIIFoldNN: Peptide:MHC Class II Binding Prediction

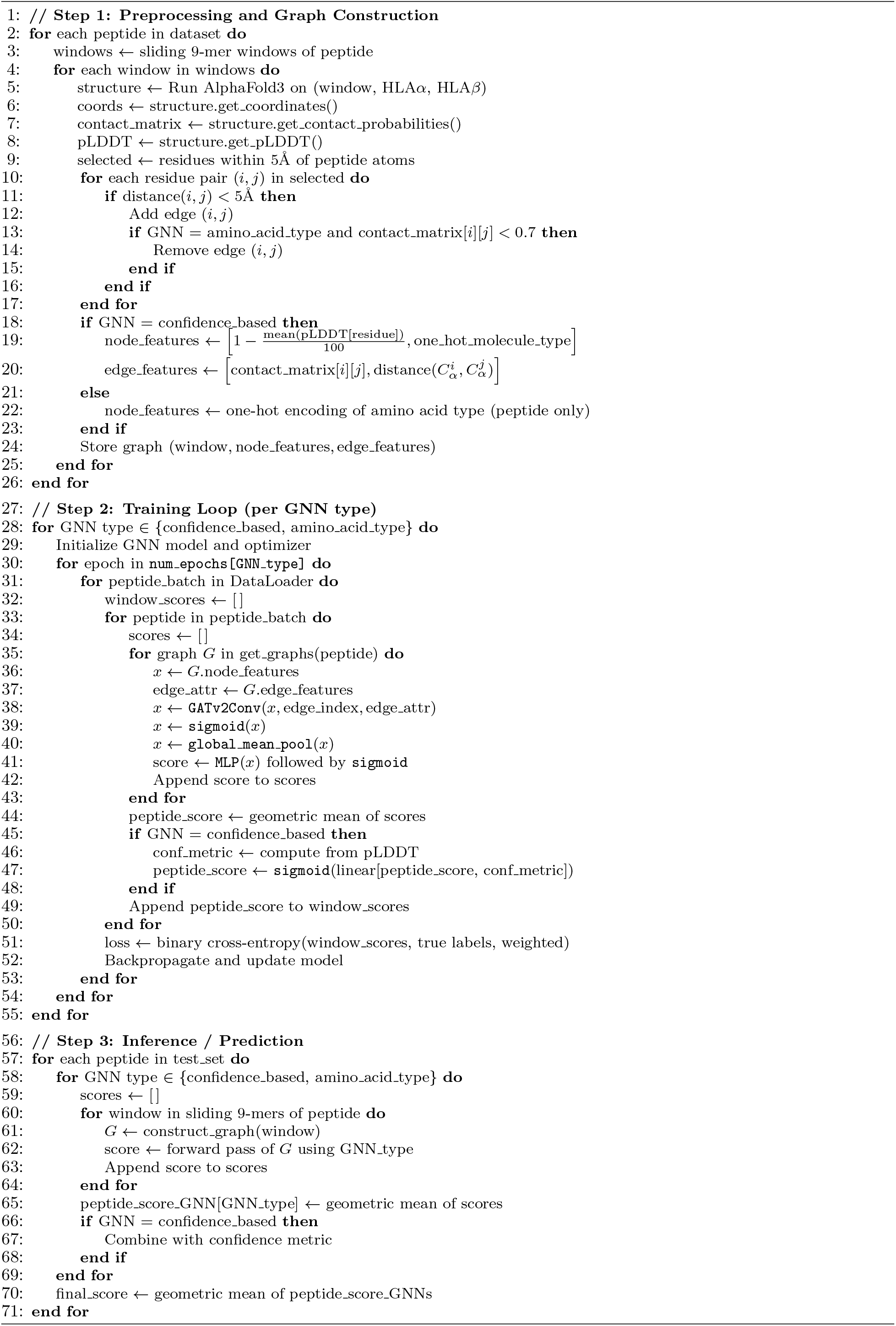

